# PROTHON: A Local Order Parameter-Based Method for Efficient Comparison of Protein Ensembles

**DOI:** 10.1101/2023.04.11.536474

**Authors:** Adekunle Aina, Shawn C.C. Hsueh, Steven S. Plotkin

## Abstract

The comparison of protein conformational ensembles is of central importance in structural biology. However, there are few computational methods for ensemble comparison, and those that are readily available, such as ENCORE, utilize methods that are sufficiently computationally expensive to be prohibitive for large ensembles. Here, a new method is presented for efficient representation and comparison of protein conformational ensembles. The method is based on the representation of a protein ensemble as a vector of probability distribution functions (pdfs), with each pdf representing the distribution of a local structural property such as the number of contacts between C*_β_* atoms. Dissimilarity between two conformational ensembles is quantified by the Jensen Shannon distance between the corresponding set of probability distribution functions. The method is validated for conformational ensembles generated by molecular dynamics simulations of ubiquitin, as well as experimentally derived conformational ensembles of a 130 amino acid truncated form of human tau protein. In the ubiquitin ensemble dataset, the method was up to 88 times faster than the existing ENCORE software, while simultaneously utilizing 48 times fewer computing cores. We make the method available as a Python package, called PROTHON, and provide a GitHub page with the Python source code at https://github.com/PlotkinLab/Prothon.

## Introduction

One of the fundamental principles that has influenced protein science is the structure-function paradigm: The idea that, for proteins with a reliable and stable three-dimensional (3D) structure, the fold invariably determines the protein’s functions.^1–3^ Protein structures are central to molecular biology and are used for example in synthetic biology^4–7^ and in drug design.^6, 8, 9^

Comparing protein structures is a common and useful practice in structural biology for the understanding of functional and evolutionary relationships.^1–3, 10–12^ Consequently, several quantitative measures for comparing protein structures have been developed; the most commonly used dissimilarity measure is the distance-based global root mean square deviation (RMSD) of atomic positions. This has however been shown to be one of the least representative of the degree of structural dissimilarity compared to contact-based methods, such as C*_β_*-C*_β_* pairwise distances, that at least in some cases are more robust and relevant.^13^ Other limitations, such as the dependence of RMSD on the accuracy of the superposition of protein structures, and its sensitivity to protein length and particularly the presence of flexible regions, have motivated more accurate measures used in Critical Assessment of techniques for protein Structure Prediction (CASP) competitions. Here metrics such as the global distance test total score (GDT-TS) and template modeling score (TM-score) are commonly used to quantify the similarity between computationally predicted structures and experimentally determined structures.^13–16^ In estimating kinetic proximity of partially disordered protein structures to the native state, generalizations of geometrical Euclidean distance have been shown to be most accurate metrics of proximity between two structures.^17, 18^

Proteins are dynamic systems, and explore a vast conformational space. Thus, comparisons between individual structures often need to be generalized to comparisons between ensembles of structures to accurately characterize macrostates of proteins. Unfolded states, partially unfolded states, or intrinsically disordered proteins (IDPs) require an ensemble description to properly characterize them. IDPs do not have a well-defined native structure, but exist as an equilibrium ensemble of diverse conformations, which inter-convert rapidly.^19–24^ Comparisons between ensembles of structures have thus been developed to treat these systems.

Unlike protein structure comparison, for which many metrics are available, efficient methods and software for comparing protein structural ensembles are much less common. The first attempt to quantitatively measure the dissimilarity between structural ensembles was an extension of the global RMSD.^25^ A different approach was proposed by Larsen and Ferkinghoff-Borg,^26^ which involves the estimation of the underlying probability distributions of protein conformational ensembles and the quantification of the overlap between these probability distributions using a symmetrized form of the Kullback-Leibler divergence. ^26–28^ Three methods were proposed for estimating the underlying probability densities, including the quasi-harmonic approximation, conformational clustering, and dimensionality reduction;^26^ all three methods had serious limitations pointed out by the authors. While the quasiharmonic approximation method is relatively fast, it is only effective for ensembles that can be described by a multivariate normal distribution. The other two methods require the calculation of all pairwise global RMSD values, a calculation scaling as ^1^ (*M*_1_ + *M*_2_)(*M*_1_ + *M*_2_ + 1), where *M*_1_ and *M*_2_ are the two ensemble sizes to be compared, which can be prohibitively expensive for large ensembles, due to the optimization of the structural superposition of structures that is required. For example, using current computing power on the Digital Research Alliance of Canada Cedar computing cluster (https://alliancecan.ca), to compare two ensembles of a peptide of 76 amino acids, each of 15000 structures, takes 58 hours (2 days and 10 hours) on a single core, and 7 hours on 48 cores running in parallel.

On the other hand, large ensembles are now routinely generated from molecular dynamics (MD) simulations, and it is often useful to compare protein ensembles generated from MD simulations using different force fields, simulation parameters, solvent conditions, or amino acid sequence. For these practical situations involving large conformational ensembles, there is a need for an effective quantitative measure and software for comparing structural ensembles, which does not require the computationally expensive process of structural superposition.

To address the problem of the high computational cost required for optimal structural alignment, as well as the potential issues of accuracy mentioned above that are associated with the use of the global RMSD of atomic positions as a structural dissimilarity measure, contact-based measures and measures utilizing internal coordinates, for example Ramachandran torsion angles, for structural comparison have previously been proposed.^13, 29^ More recently, a measure for comparing IDP ensembles based on C*_α_*-C*_α_* distance matrices was described in Ref.^30^ Although the method described in reference^30^ is superposition-independent, therefore requiring relatively lower computational cost, it only considers distance distribution averages rather than the full distributions themselves.

In the present work, we introduce and implement a new generalized method for the efficient representation and comparison of protein conformational ensembles. The method involves the utilization of the Jensen Shannon Distance (JSD) metric for the quantification of the difference between probability distribution functions representing the distributions of a local structural property *θ* of polypeptide chains that constitute the protein ensemble under consideration. A related approach for comparing conformational ensembles was previously introduced by McClendon, Hua, Barreiro, and Jacobson, in which the Kullback-Leibler Divergence was used to quantify the differences between the distributions of the Ramachandran torsion angles.^31^ In principle, any local structural property can be used in our method. Local structural properties including per residue solvent accessible surface area (SASA),^32, 33^ virtual C*_α_*-C*_α_* bond angle (CABA) and torsion angle (CATA),^34^ C*_α_* contact number (CACN), and C*_β_* contact number (CBCN)^35, 36^ have been implemented in other contexts, and could be applied to the method introduced here. In this manuscript we use CBCN. The method of protein ensemble comparison described here was implemented using the Python programming language, and is made available as a Python package, called PROTHON (PROtein *θ* comparisON), for easy access to the computational structural biology scientific community. PROTHON provides an easy-to-use protein ensemble comparison program and a simple code framework for extension to include other local structural properties. We provide a GitHub page with the Python source code at https://github.com/PlotkinLab/Prothon.

## Description

In this section, we describe the algorithm for the efficient representation and comparison of protein conformational ensembles.

### Matrix Representation of a Protein Ensemble

Consider a protein ensemble with *M* conformations and *N* local structural property values (e.g. CBCN), as described above. Typically *N* is proportional to the number of amino acids in the chain. Let *θ* be a local structural property, such as the per-residue C*_β_* contact number (CBCN)^35, 36^ (see section on C*_β_* Contact Number Local Structural Property). In a given conformation *m* of the polypeptide chain, there are *N θ* values, and for a given local index such as a given residue, there are *M θ* values. Thus *θ* is a function of (*m, n*), the conformation index and the residue index. As parameterized by the local property *θ*, the full ensemble of conformations can be represented by an *M* × *N* matrix, X, with elements *X_m,n_* given by Eq. 1:

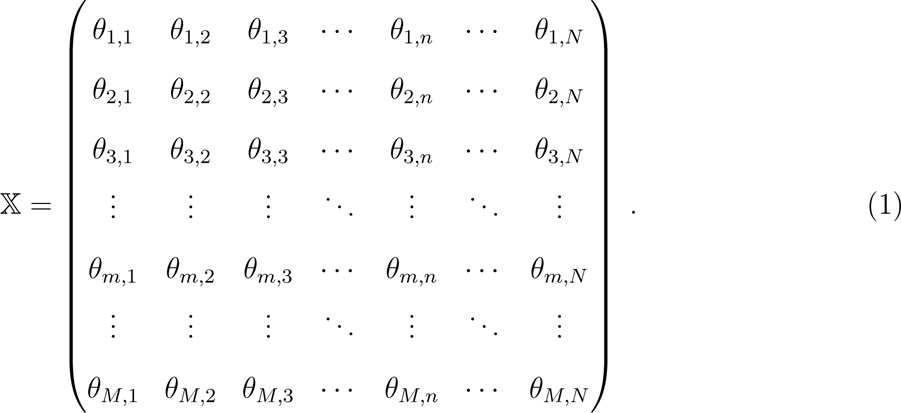

Each row of matrix 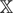 can be thought of as a vector representing the *θ* values in a conformation, e.g. 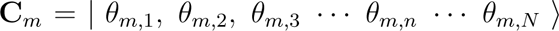, and each column is an ensemble 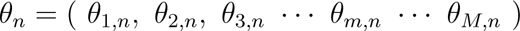 of local values of *θ* across all conformations in the protein ensemble. A typical size of 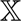 might be a (*M, N*)=10,000 × 100 dimensional matrix.

### Protein Ensemble Represented as a Vector of Local Ensembles

The above matrix representing a protein ensemble, as described above in Matrix Representation of a Protein Ensemble, can alternatively be written as a vector of local ensembles *θ_n_*(see Eq. 2).

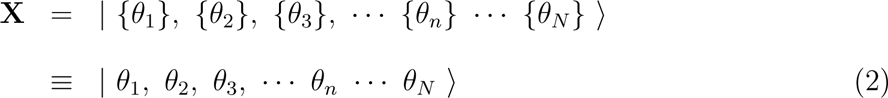

Each *ensemble θ_n_* can further be represented by a probability distribution function (pdf),e.g. the ensemble of *θ*_1_ can be represented as *p*(*θ*_1_) ≡ *p*_1_. The conformational ensemble of the whole protein may then be written as

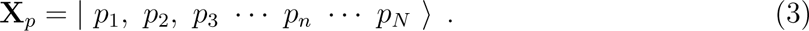

In the present work, we represent the pdfs in Eq. (3) by one-dimensional (1D) distribution functions. The probability distribution function *p_n_* representing each *θ_n_* is determined by Gaussian kernel-density estimation with the Silverman bandwidth estimator method,^37^ which is implemented here using SciPy. ^38^ A conventional histogram with suitable choice of binning may be used as well, however we choose Gaussian kernel-density estimation here because of its advantages of a smoother estimate for the density distribution, which is convenient for ensemble comparison. To implement the ensemble comparison analysis below, we compare two (or more) distributions 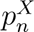, 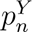 for the same *θ_n_* as follows. The minimum and maximum sampled values of *θ_n_* are first determined, then the values of 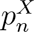 and 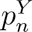 at *ν* = 100 equally-spaced points between *θ_min_* and *θ_max_* are obtained. Each distribution is represented by *ν* = 100 values:

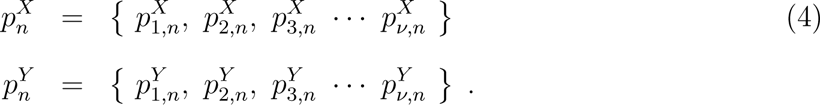

The value *ν* may be increased beyond 100 if necessary. The choice of *θ_min_* and *θ_max_* typically results in one or more of the distributions having values of zero near one or both of the limits of *θ*.

### Quantifying Similarity Between Protein Ensembles

Here, we propose an efficient and effective dissimilarity measure for protein conformational ensembles, by comparing pdfs representing a local property distribution using the Jensen Shannon Distance (JSD) metric. The JSD is an information theory-based measure for comparing two probability distributions. A related non-metric version of JSD, the Jensen Shannon Divergence, has been used to quantify the dissimilarity between protein ensembles in a previous study. ^28^ Employing this procedure can require high computational cost however, and may have limited practical applicability to large ensembles.

The ensemble representation as described in Protein Ensemble Represented as a Vector of Local Ensembles above allows for the calculation of both local and global similarity between two protein ensembles. Given two protein ensembles 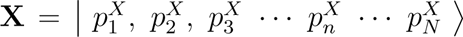 and 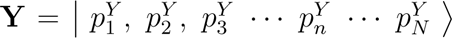 which are represented as vectors of pdfs (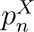 and 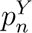) of a local structural property, the dissimilarity between ensembles **X** and **Y** is estimated as the *N*-dimensional vector, termed here as the dissimilarity vector, given by:

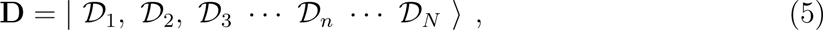

where each component of **D**, 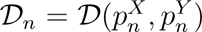, is the JSD between the probability distributions representing the corresponding local structural property values *θ_n_* in the polypeptide chain in the ensembles **X** and **Y**. The JSD is defined as

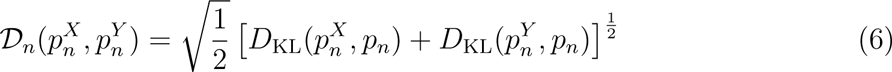

where

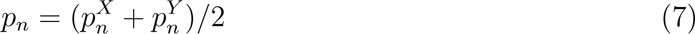

and e.g

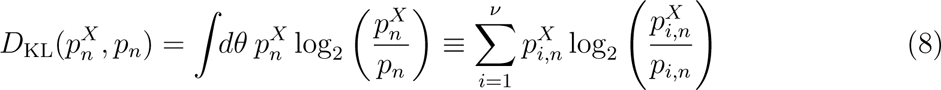

is the Kullback-Leibler divergence between probability distributions 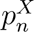 and *p_n_*.^27^ 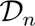 has a lower and upper bound of 0 and 1 respectively. A value of 0 indicates that two ensembles are identical while a value of 1 suggests that they are completely different.

The global dissimilarity *D* between ensembles **X** and **Y** is calculated as the average over the *N* local dissimilarities.

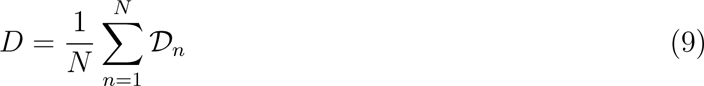

Motif dissimilarity between ensembles can also be determined in a similar fashion by averaging over local structural property values belonging to a particular motif. The dissimilarity vector in Eq. (5) allows ensemble comparisons for proteins with both structured and disordered domains.

### C***_β_*** Contact Number Local Structural Property

The C*_β_* contact number (CBCN) implemented in PROTHON is defined by the number of C*_β_* atoms on other residues separated by 3 or more amino acids from a given residue, that are located within a sphere of radius *r_o_* centred on the C*_β_* atom of the residue of interest. Here we implement this criterion with a smooth cutoff function:

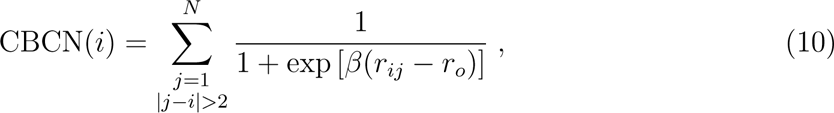

where the sum runs over the *N* C*_β_* atoms *j* that may be in contact with C*_β_* atom *i* such that |*j* − *i*| *>* 2, and *r_ij_*is the distance between C*_β_* atoms *i* and *j*, belonging to residues *a* and *b* with a sequence separation |*a* − *b*| *>* 2. The parameters *β* and *r_o_* were taken to be 50 nm*^−^*^1^ and 1 nm, respectively. A similar method for calculating contact number was previously used by Kinjo, Horimoto, and Nishikawa^35^ and Yuan.^36^

### Measuring Significant Dissimilarity

To compare two protein ensembles 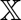 and 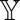 with *M_x_* and *M_y_* number of conformations, respectively, we consider only local dissimilarities (i.e. 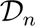) between the ensembles that are statistically significant relative to dissimilarities within each ensemble. 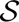 samples of each ensemble are generated by randomly sampling (with replacement) 1000 conformations from ensembles 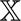 and 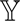, respectively. The local dissimilarities 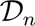 are then calculated between the 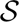 samples within the same ensemble and across different ensembles, which results in 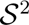 inter-ensemble sample dissimilarities, and 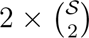 intra-ensemble sample dissimilarities. In this present work, we take 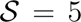 = 5 samples, so that there are 25 inter-ensemble sample dissimilarities and 20 intra-ensemble sample dissimilarities (10 for 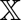 and 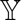, respectively).

The statistical significance of local dissimilarities 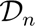 are calculated using the Mann–Whitney U test,^38–40^ which is a non-parametric test of the null hypothesis: the probability of *x* being greater than *y* is equal to the probability of *y* being greater than *x* for randomly selected values *x* and *y* from two populations. Only dissimilarities 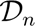 with *p*-values *<* 0.05 are taken to be statistically significant.

## Validation

The validity and computational efficiency of the introduced method for the comparison of protein ensembles implemented in PROTHON is demonstrated in the sections below with computationally generated, experimentally derived, structured, and intrinsically disordered protein ensembles.

### Comparing Computationally Generated Ensembles of Ubiquitin

Ubiquitin is a regulatory protein that exists in all eukaryotic cells. It performs various functions through conjugation to many different target proteins.^41^ Ubiquitin consists of 76 amino acids and has a molecular mass of about 8.6 kDa.

To generate conformational ensembles for the natively structured protein ubiquitin, we used the 3D coordinates of its atoms downloaded from the protein data bank (PDB ID: 1UBQ^42^) as the initial state to perform molecular dynamics (MD) simulations with the CHARMM36m force field^43^ using GROMACS (version 2019.2).^44^ All simulations were performed in explicit solvent (TIP3P water model, and 150 mM NaCl) at 300 K and 1 bar with periodic boundary conditions. Five independent MD runs were performed for the folded state, and 5 independent MD runs were performed for 5 partially folded states of ubiquitin. Each MD run was 50 ns. The partially folded states were obtained by performing global unfolding MD simulations, using PLUMED,^45^ along a collective coordinate *Q* defining the fraction of native contacts:

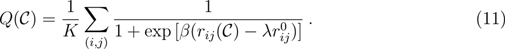

In (Eq. 11), the sum runs over *K* pairs of native contacts (*i, j*), where *K* is determined from all atom pairs within 0.45 nm in the native structure. 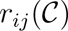 is the distance between atoms *i* and *j* in conformation 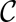, belonging to residues *a* and *b* with a sequence separation |*a*−*b*| *>* 3, and 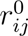 is the native distance between atoms *i* and *j*. The parameters *β* and *λ* were taken to be 50 nm*^−^*^1^ and 1.5 respectively. A similar method for defining fraction of native contacts has been used in several previous studies, see e.g. references 46,47.

A time-dependent harmonic potential *V* (*Q, t*)

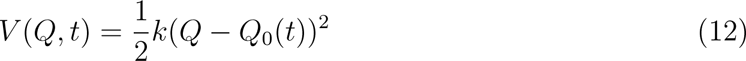

was applied to move the center of the bias linearly from *Q*_0_ = 1 to *Q*_0_ = 0.95, 0.90, 0.85, 0.80, 0.75, respectively for 5 partially folded states. The global unfolding bias was applied for 10 ns and then held fixed for 40 ns. The spring constant *k* in Eq. (12) was taken to be 10^7^ kJ·mol*^−^*^1^.

Conformations were sampled at equal time intervals of 40 ps from the last 40 ns of five independent simulations for each of the above values of *Q*_0_, giving a total of 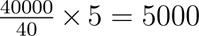 conformations in each of the folded and partially folded ensembles. Fig. 1 shows randomly-selected structures from the folded ensemble and for five partially folded ensembles that correspond to the five *Q*_0_ values.

**Figure 1:**
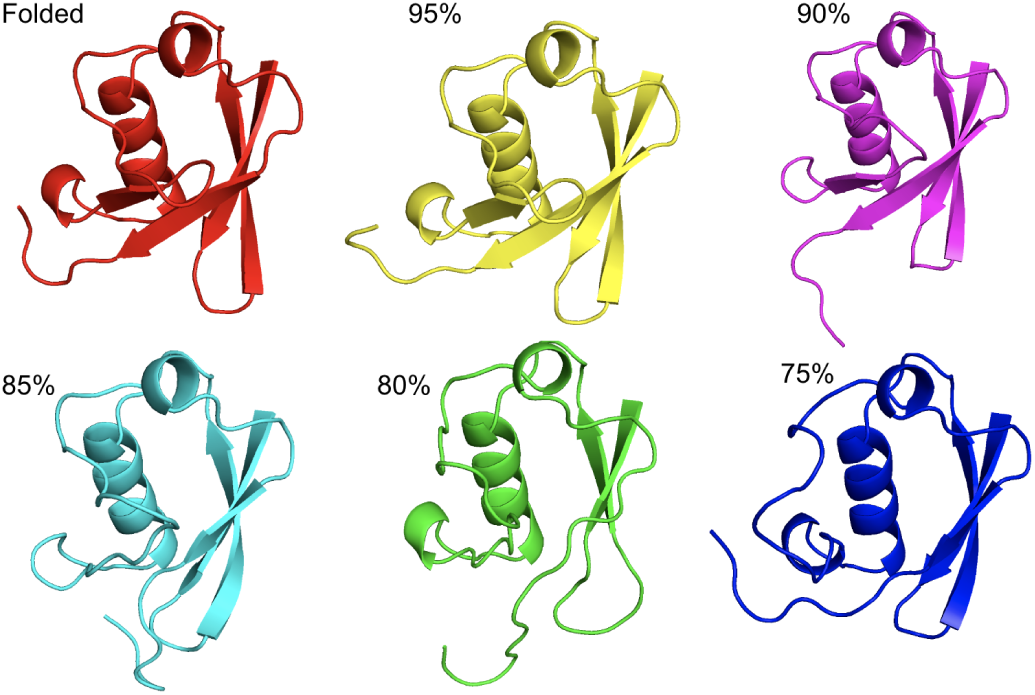
Representative structures from ubiquitin ensembles. The folded **(red)** and partially folded: 95% folded **(yellow)**, 90% folded **(magenta**), 85% folded **(cyan)**, 80% folded **(green)**, and 75% folded **(blue)** ubiquitin ensembles. All images were rendered with PyMOL molecular visualization system (https://pymol.org).

The degree of nativeness (or foldedness) of ubiquitin was taken as the fraction of native contacts *Q*. To visualize these ensembles, we represented them as CBCN *M* × *N* matrices (see Matrix Representation of a Protein Ensemble), and as pairwise RMSD *M* × *M* matrices (as used in ENCORE^28^), where *M* = 30000 is the total number of structures and *N* = 70 is the number of C*_β_* atoms in each structure. The high dimensional CBCN and RMSD matrices were then reduced to 2-dimensions (2D) using principal component analysis (PCA^48^), multi-dimensional scaling (MDS^49^), and t-distributed stochastic neighbor embedding (t-SNE^50^) dimensionality reduction methods (see Fig. 2). PCA, MDS, and t-SNE were implemented using Scikit-learn.^51^

**Figure 2:**
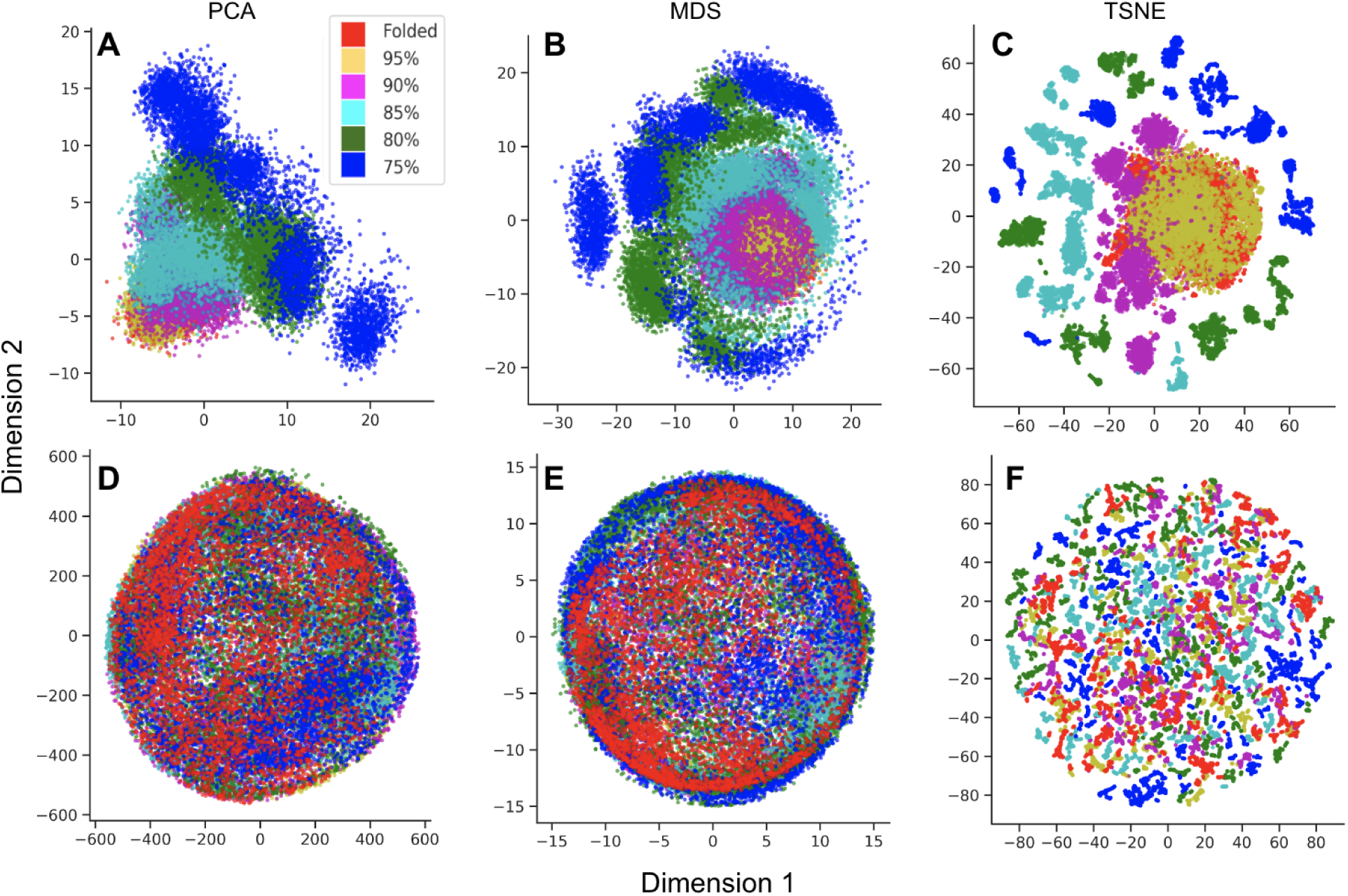
Representation and dimensionality reduction of ubiquitin ensembles. The folded (red) and partially folded (yellow: 95% folded, magenta: 90% folded, cyan: 85% folded, green: 80% folded, and blue: 75% folded ubiquitin ensembles are represented using the PROTHON C*_β_* contact number (CBCN) matrix **(top panels A, B, C)** and ENCORE RMSD matrix **(bottom panels D, E, F)**. Dimensionality reduction to 2D using **(left)** principal component analysis (PCA) for **(A)** CBCN and **(D)** RMSD matrix representations, **(middle)** multi-dimensional scaling (MDS) for **(B)** CBCN and **(E)** RMSD matrix representations, and **(right)** t-distributed stochastic neighbor embedding (t-SNE) for **(C)** CBCN and **(F)** RMSD matrix representations.

In Fig. 2, the ensembles are color coded as folded (red) and partially folded (yellow: 95% folded, magenta: 90% folded, cyan: 85% folded, green: 80% folded, and blue: 75% folded). At this 2D level of resolution, whereby the PCA explained variance is ≈ 40%, the ensembles are more readily distinguishable for the CBCN matrix representation (Fig. 2 panels A, B, and C) than the pairwise RMSD matrix representation (Fig. 2 panels D, E, and F). As expected for these ubiquitin ensembles, the proximity in the projected space of structures in the partially folded ensembles to structures in the natively folded ensemble decreases with decreasing degree of foldedness (e.g. yellow data points are closer to the red than blues are to the red), and the degree of spread increases with decreasing degree of foldedness (i.e. there is more spatial variation in the projected spaces for blue than there is for yellow.) (see Fig. 2 top panels A,B,C). These observed relationships between the ensembles using the CBCN matrix are visually absent when using the RMSD matrix representation (see Fig. 2 bottom panels D,E,F).

The distinct clusters of the ensemble with 75% natively folded ensemble (e.g. blue in Fig. 2 A), correspond to the different independent simulations (see Fig. S1). To enhance clarity and interpretation of the data, each ensemble in Fig. 2 is plotted separately (non-overlapping) in Figs. S2-S7.

Fig. 3A shows the dissimilarity of partially folded ensembles of ubiquitin from the natively folded ensemble at different degrees of partial nativeness, 75% *< Q <* 95%, quantified using the C*_β_* contact number (CBCN) local structural property as implemented in PROTHON, and the global RMSD structural property as implemented in ENCORE, both using the PCA dimensionality reduction method.^28^ Both the CBCN local structural properties and the global RMSD structural property are able to distinguish the partially folded ensembles from the folded ensemble and correctly determine the order of nativeness without prior knowledge of how the ensembles were obtained (Fig. 3A).

**Figure 3:**
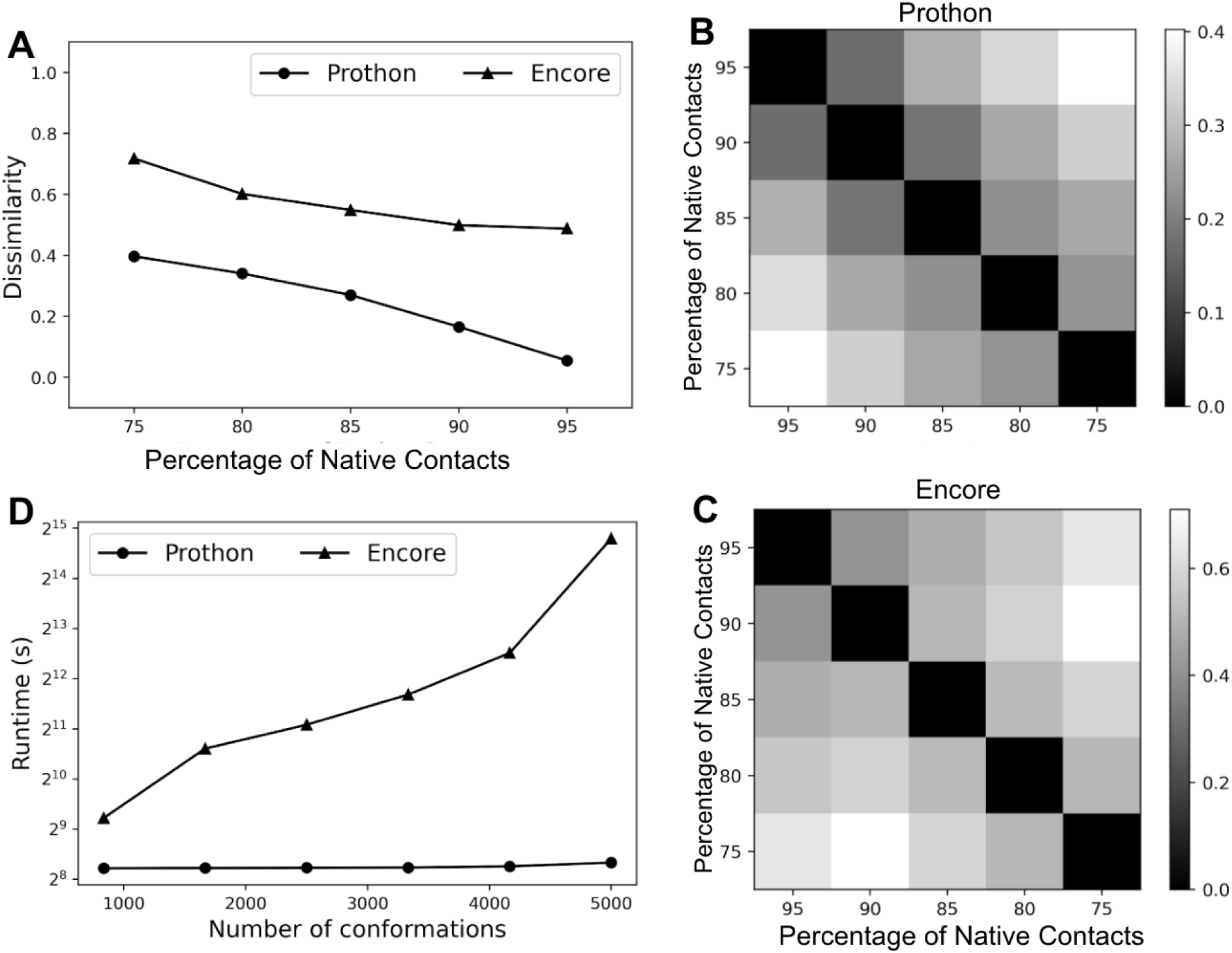
Comparing partially folded ensembles of ubiquitin. **(A)** Dissimilarity between partially folded ensembles (75% *< Q <* 95%) and the folded equilibrium ensemble of ubiquitin. Dissimilarity is quantified using the local structural property of C*_β_* contact number (circles) implemented in PROTHON, and RMSD similarity is implemented in ENCORE (triangles). **(B)** Pairwise dissimilarity between the 5 partially folded ensembles of ubiquitin in panel (A) using PROTHON. **(C)** Pairwise dissimilarity between the 5 partially folded ensembles of ubiquitin in panel (A) using ENCORE.^28^ **(D)** Wall-clock time (computational efficiency) of PROTHON compared with ENCORE. ENCORE calculations were run on 48 cores; PROTHON calculations were run on a single core.

We also examine four additional local structural properties to demonstrate the generality of the PROTHON approach for ensemble comparison: the C*_α_* contact number (CACN), virtual C*_α_*-C*_α_* bond angle (CABA) and torsion angle (CATA),^34^ and per-residue solvent accessible surface area (SASA).^32, 33^ Fig. S8 plots the dissimilarity *vs.* the percentage of native contacts, indicating that all local structural properties accurately distinguish between the partially folded ensemble and the folded ensemble of Ubiquitin. The order parameter we focus on here, CBCN, has among the largest relative differences between the ensembles, and does not exhibit the same degree of non-linear “plateauing” as some of the other order parameters do.

Fig. 3 panels B and C show a heatmap greyscale-coded matrix of pairwise dissimilarity between the 5 partially folded ensembles of ubiquitin, as quantified using CBCN and RMSD respectively. As expected, the dissimilarity increases (becomes lighter in the matrices) as one moves away from the diagonal in all 4 directions (up, down, left, and right). The dissimilarity is zero when comparing an ensemble with itself (black diagonal) The dissimilarity increases as one moves away from the diagonal when using RMSD as a dissimilarity measure, but an exception is noticeable: The ensemble with 75% native contacts is more dissimilar to the ensemble with 90% native contacts than it is to the ensemble with 95% native contacts; this nonmonotonicity is unexpected. These results show that the methods of protein ensemble representation and comparison as implemented in PROTHON may be more effective than those using RMSD.

### Comparing Experimentally Derived Ensembles of the Tau K18 Domain

Tau is an intrinsically disordered protein (IDP), whose misfolding and aggregation is implicated in many neurodegenerative diseases including Alzheimer’s disease (AD), Pick’s disease (PiD), chronic traumatic encephalopathy (CTE), corticobasal degeneration (CBD), and progressive supranuclear palsy (PSP).^52–54^ Tau K18 domain (tauK18) is a 130-residue truncated human tau protein consisting of the four microtuble binding repeats of tau. Five ensembles of tauK18 were download from the Protein Ensemble Database (PED ID: PED00017).^55^ These ensembles represent different models of tauK18 that were derived by first sampling random coil conformations and then selecting those conformations that fit experimentally determined chemical shifts and residual dipolar couplings in NMR data.^56^

Table 1 shows the pairwise conformational dissimilarity of the 5 tauK18 ensembles. Conformational dissimilarity here was quantified using the C*_β_* contact number (CBCN) local structural property implemented in PROTHON. The dissimilarity values are all small (average dissimilarity ≈ 0.03), suggesting that all 5 generated ensembles represent similar ensembles, in agreement with the result obtained by Lazar *et al.*.^30^

**Table 1:**
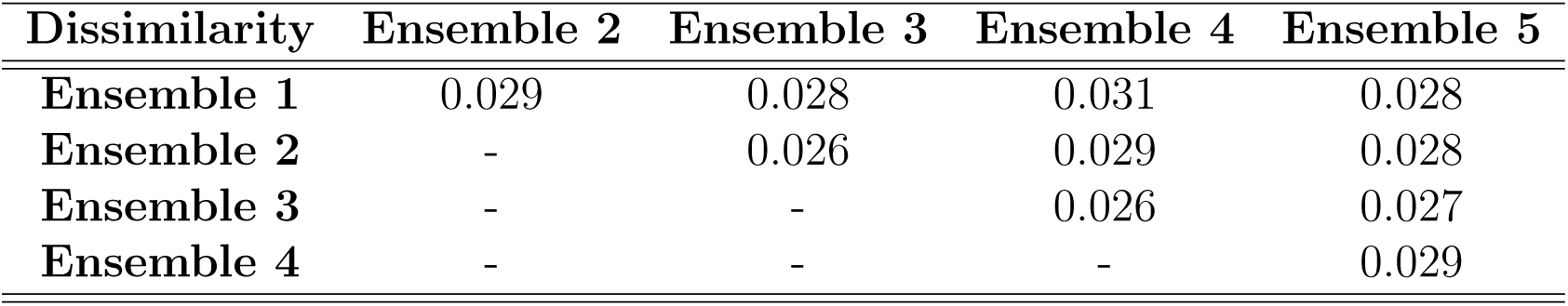
Comparing intrinsically disordered ensembles of tau K18. The pairwise dissimilarity between 5 ensembles of tau K18 domain quantified using the C_β_ contact number local structural property implemented in PROTHON.

| Dissimilarity | Ensemble 2 | Ensemble 3 | Ensemble 4 | Ensemble 5 |
| --- | --- | --- | --- | --- |
| **Ensemble 1** | 0.029 | 0.028 | 0.031 | 0.028 |
| **Ensemble 2** | - | 0.026 | 0.029 | 0.028 |
| **Ensemble 3** | - | - | 0.026 | 0.027 |
| **Ensemble 4** | - | - | - | 0.029 |

### Computational Efficiency

To evaluate the computational efficiency of PROTHON, we recalculated the pairwise dissimilarity between the 5 ubiquitin partially folded ensembles described in Comparing Computationally Generated Ensembles of Ubiquitin. The calculation of the pairwise ensemble dissimilarity was repeated 6 times using PROTHON, and also using the publicly available program ENCORE.^28^ For the *i*th run, where *i*, 1 ≤ *i* ≤ 6, a subset of conformations were selected from the 5000 total conformations that make up each ensemble by choosing conformations at *i* regular frame intervals, for a total of 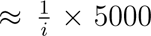 conformations in each ensemble, i.e. every *i*th frame is sampled, for 1 ≤ *i* ≤ 6. Since PROTHON is currently a serial code, each run used a single core on a 2020 MacBook Pro, while 48 cores were available for ENCORE, which has parallel code, on the Digital Research Alliance of Canada Cedar computing cluster (https://alliancecan.ca).

Fig. 3D shows the wall clock time required to calculate the pairwise ensemble dissimilarity for the 5 ubiquitin partially folded ensembles, with varying number of conformations in each ensemble using PROTHON and ENCORE, respectively. That is, we performed 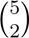 ensemble dissimilarity calculations to calculate 10 values of D in Eq. (9) for PROTHON and 10 values of the Jensen Shannon divergence for ENCORE. While the time required to run ENCORE increases exponentially with the number of conformations (the y-axis of the plot is on a log scale), the time required to run PROTHON essentially remains the same at ≈ 300 seconds = 5 minutes. We anticipate that for large ensemble sizes the scaling should be linear in the number of conformations (times the chain length), because filling in the matrix 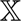 in Eq. (1) is sufficient to determine the distribution **X***_p_* in Eq. (3). When *i* = 6 and each ensemble had 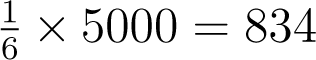 conformations, it takes ENCORE 596 seconds = 9 minutes, 56 seconds, which is twice that of PROTHON (which takes 298 seconds = 4 minutes, 58 seconds), to run. When *i* = 1 and each ensemble had 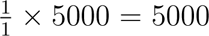 conformations, the time it takes ENCORE to run increases dramatically to 28,406 seconds = 7 hours, 53 minutes, 26 seconds, which is more than 88 times that of PROTHON (which takes 322 seconds = 5 minutes, 22 seconds) despite ENCORE using 48× the cores of PROTHON. If ENCORE runs on a single core, the corresponding calculation time for 5000 conformations is 58 hours (2 days and 10 hours), which would be 2^17.7^ in Fig. 3D. We also note that this is not linear scaling for ENCORE; the 48 core calculation shows a speedup of only 7.4×.

Although it may be possible to achieve speed-up in practice by substituting the RMSD matrix used in ENCORE with a matrix using an alignment-free structural comparison, such as dRMSD,^57–59^ fraction of common contacts (Q),^60^ local distance test,^61^ global distance test (GDT),^62^ or minimal Euclidean distance,^18^ the computational complexity would still remain at 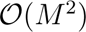, where *M* is the number of structures to be compared. This is in contrast to PROTHON, which achieves a computational complexity of 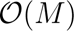. We reserve for future work a more extensive, systematic comparison between the metric we propose here and other metrics proposed in the literature.

## Software and Data Availability

PROTHON is freely available. Users can run PROTHON by downloading the Python file Prothon.py from github under the GPLv3 license at https://github.com/PlotkinLab/ Prothon. The snippet below shows how to use PROTHON to calculate ensemble dissimilarity in only 12 lines of Python code. The code below was used to generate the data for the plot in Fig. 3A; a description follows.

~~~
1 **from** Prothon **import** Prothon
2 **import** numpy as np
3
4 data = ['Q99 . dcd','Q75 . dcd','Q80 . dcd','Q85 . dcd','Q90 . dcd','Q95 . dcd']
5 topology = 'topology . pdb'
6
7 prothon = Prothon ( data = data, topology = topology )
8 ensembles = prothon . ensemble_representation ( measure = 'CBCN')
9 x_min, x_max = (np. **min**( ensembles), np. **max**( ensembles))
10
11 dissimilarity = []
12
13 **for** ensemble **in** ensembles [1:]:
14    d = prothon . dissimilarity ( ensemble, ensembles [0], x_min = x_min, x_max = x_max )
15    dissimilarity . append ( d [0])
16
17 **print** ( dissimilarity )
~~~

The Prothon class is first imported from the Prothon package (line 1), followed by the creation of a Prothon object (line 7) initialized with the topology in pdb format (line 5) of the conformations contained in these ensembles, and a list of the ensembles (line 4) to be compared. The dcd format has been used here for the ensemble data, but any format supported by the MDTraj package^33^ is allowed. The Prothon *ensemble representation* function is then used to represent the ensembles as described in the present work using the C*_β_* contact number (CBCN) local structural property (line 8). The minimum and maximum values of CBCN in all ensembles are obtained in line 9 using numpy, which is imported in line 2.

To calculate the dissimilarity between two ensembles, the Prothon *dissimilarity* function is used (line 14), which returns a list *d* with 3 objects: the global dissimilarity, local dissimilarity values, and the *p*-values for the local dissimilarity, (i.e statistical significance, with dissimilarity being significant if the p-value *<* 0.05). There are 70 elements for the local dissimilarity and for the *p*-values, corresponding to the number of C*_β_* atoms. The global dissimilarity between each ensemble and the first ensemble is saved (line 15) to the dissimilarity list (created in line 11), which is finally displayed in line 17.

The ubiquitin molecular dynamics-generated ensemble dataset (229 MB zip file) is available at https://doi.org/10.5281/zenodo.7792288.

## Conclusion

We have developed a new generalized method for the efficient representation and comparison of protein ensembles. Our newly developed method was implemented in the Python programming language, and made freely available as a Python package called PROTHON to the computational structural biology community. The method involves the representation of a protein ensemble as a vector of probability distribution functions of a local quantity involving each amino acid. Each probability distribution function is estimated, using Gaussian kernel density estimation, from the distribution of a local structural property of the polypeptide chain. The Jensen-Shannon distance between corresponding probability distribution functions then quantifies the dissimilarity between protein ensembles. Here, the C*_β_* contact number (CBCN) was used as a local structural property, but in principle any local structural property can be used in PROTHON. Examples include solvent accessible surface area (SASA) per residue,^32, 33^ virtual C*_α_*-C*_α_* bond angle (CABA) and torsion angle (CATA),^34^ and C*_α_* contact number (CACN). PROTHON was shown to be effective in correctly distinguishing computationally generated ensembles of ubiquitin and experimentally derived ensembles of a 130 amino acid fragment of tau protein. The computational efficiency of PROTHON, when compared to the publicly available software ENCORE,^28^ can simultaneously yield up to an 88-fold gain in wall-clock time while benefiting from a 48-fold reduction in the number of computing cores required.

## Supporting Information Available

The Supporting Information is available free of charge at https://pubs.acs.org/

1. Representation, dimensionality reduction, and comparison of ubiquitin ensembles (PDF)

## Authors

Adekunle Aina – Department of Physics and Astronomy, The University of British Columbia, Vancouver, BC, V6T 1Z1, Canada.

Shawn C.C. Hsueh – Department of Physics and Astronomy, The University of British Columbia, Vancouver, BC, V6T 1Z1, Canada.

## Author Contributions

Conceptualization: A.A.; Methodology: A.A. and S.S.P.; Software: A.A.; Data analysis: A.A. and S.H.; Visualization: A.A.; Resources: S.S.P; Writing—original draft: A.A.; Writing—review and editing: S.S.P.; Supervision: S.S.P.; Funding acquisition: S.S.P.

## Notes

The authors declare no competing financial interest.

## Acknowledgement

This research was supported by Canadian Institutes of Health Research Transitional Operating Grant 2682, Alberta Innovates Research Team Program Grant PTM13007, Compute Canada Resources for Research Groups RRG 3071, and UBC ARC Sockeye Advanced Research Computing (https://doi.org/10.14288/SOCKEYE, 2019). A.A. has received support from a UBC Four-Year Fellowship. S.C.C.H. has received support from a NSERC CREATE-Ecosystem Services, Commercialization, and Entrepreneurship (ECOSCOPE) scholarship.

## Supplementary Figures

**Fig.S1:**
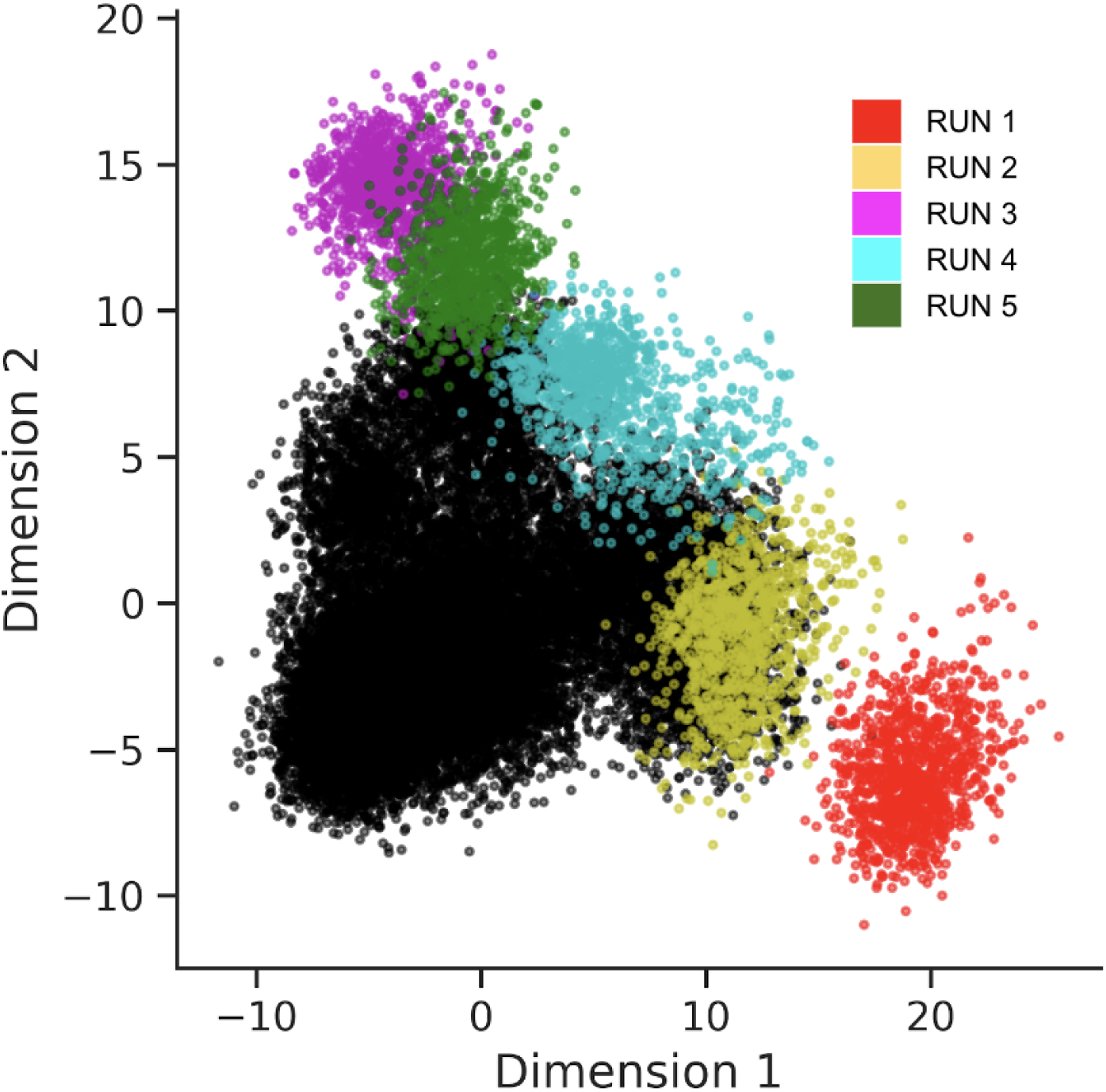
Representation and dimensionality reduction of ubiquitin ensembles. The 5 independent simulation runs of the 75% folded ubiquitin ensemble are represented in 5 colors: **(red)** first run, **(yellow)** second run, **(magenta)** third run, **(cyan)** fourth run, and **(green)** fifth run. All other ensembles are shown in **black**. Ensembles are represented using the PROTHON C*_β_* contact number (CBCN) matrix. Dimensionality reduction to 2D was performed using principal component analysis (PCA).

**Fig.S2:**
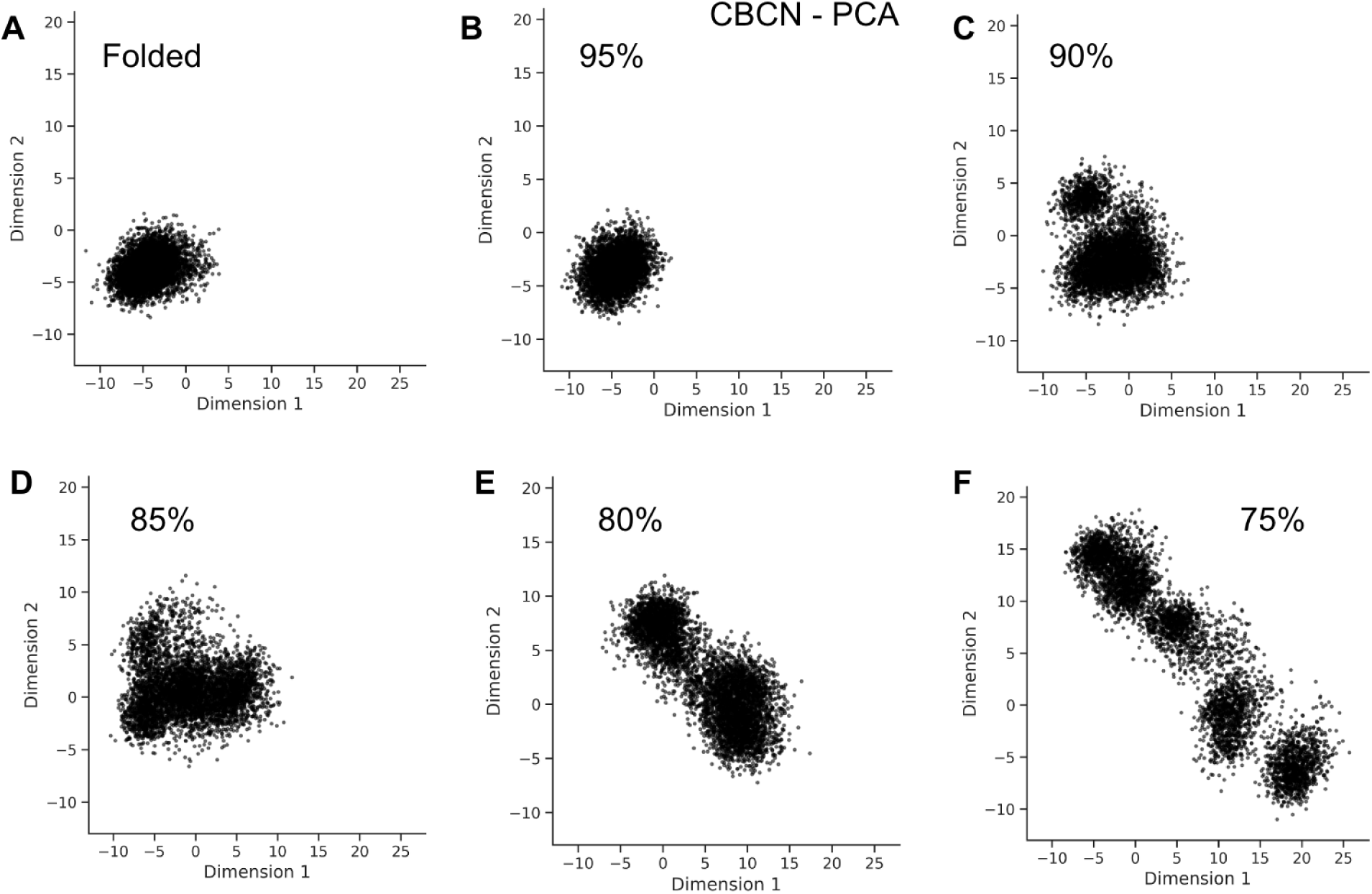
Representation and dimensionality reduction of ubiquitin ensembles. **(A)** The folded and partially folded **(B)** 95% folded, **(C)** 90% folded, **(D)** 85% folded, **(E)** 80% folded, and **(F)** 75% folded ubiquitin ensembles are represented using the PROTHON C*_β_* contact number (CBCN) matrix. Dimensionality reduction to 2D was performed using principal component analysis (PCA).

**Fig.S3:**
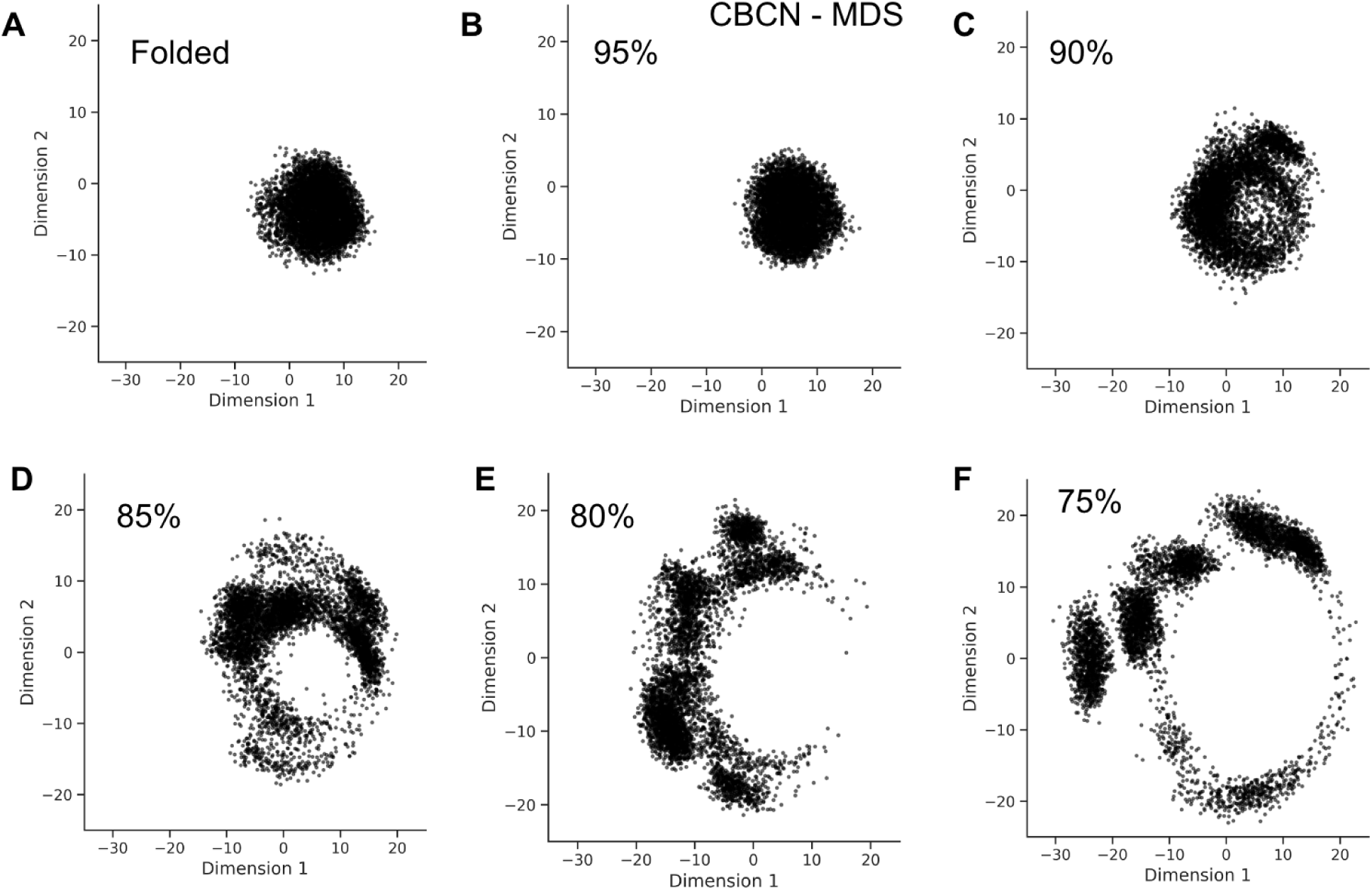
Representation and dimensionality reduction of ubiquitin ensembles. **(A)** The folded and partially folded **(B)** 95% folded, **(C)** 90% folded, **(D)** 85% folded, **(E)** 80% folded, and **(F)** 75% folded ubiquitin ensembles are represented using the PROTHON C*_β_* contact number (CBCN) matrix. Dimensionality reduction to 2D was performed using multi-dimensional scaling (MDS).

**Fig.S4:**
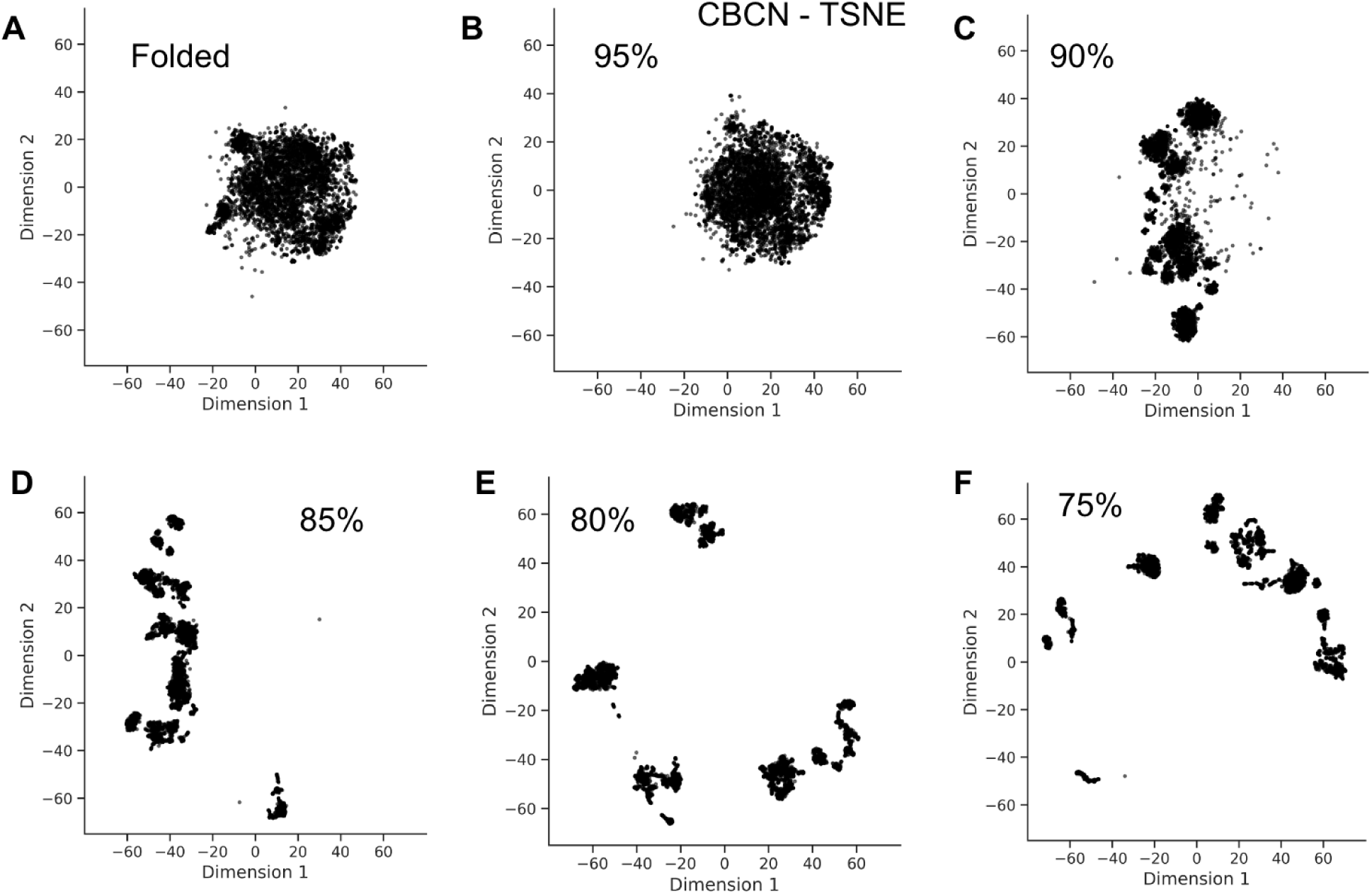
Representation and dimensionality reduction of ubiquitin ensembles. **(A)** The folded and partially folded **(B)** 95% folded, **(C)** 90% folded, **(D)** 85% folded, **(E)** 80% folded, and **(F)** 75% folded ubiquitin ensembles are represented using the PROTHON C*_β_* contact number (CBCN) matrix. Dimensionality reduction to 2D was performed using t-distributed stochastic neighbor embedding (t-SNE).

**Fig.S5:**
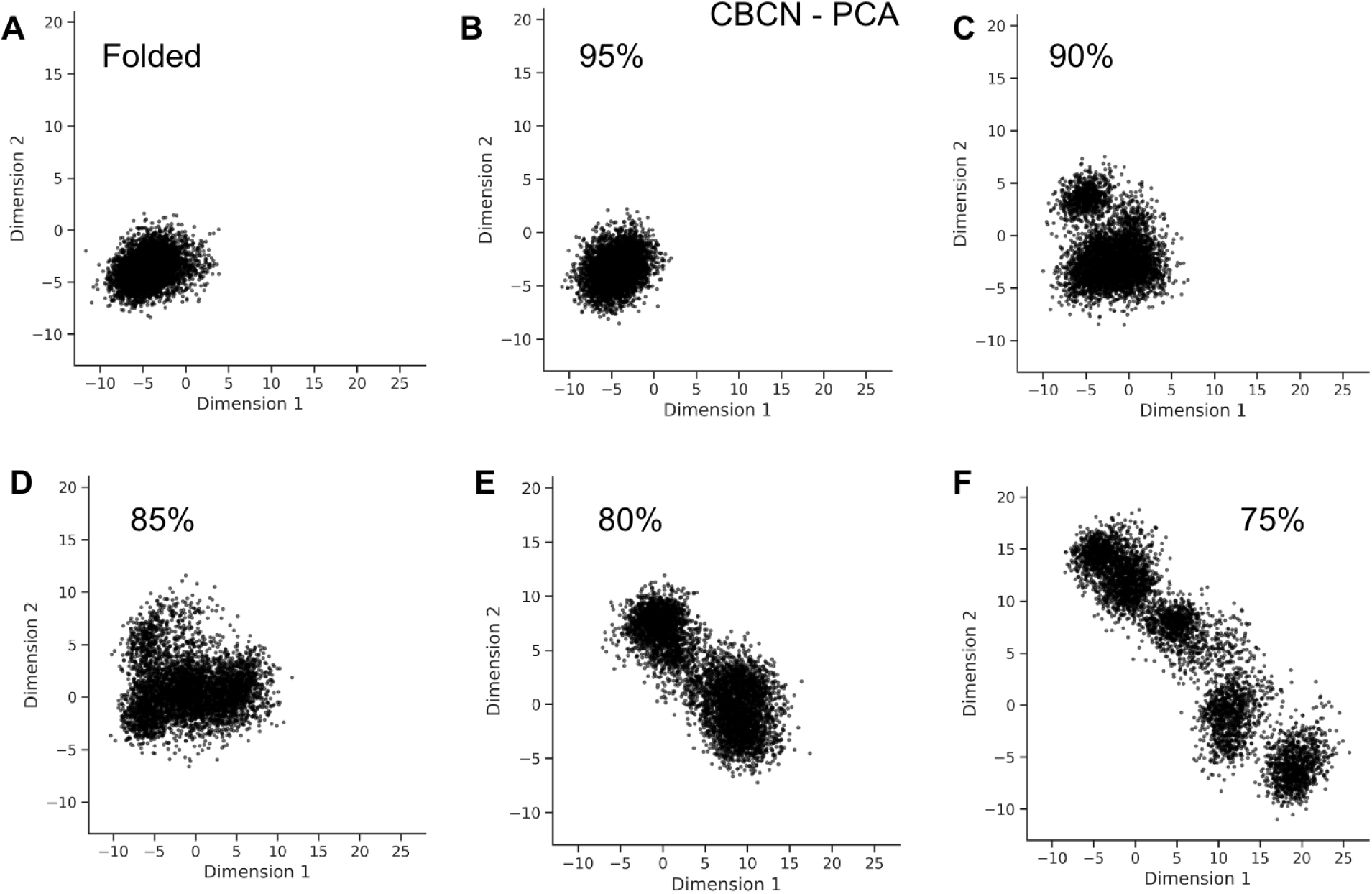
Representation and dimensionality reduction of ubiquitin ensembles. **(A)** The folded and partially folded **(B)** 95% folded, **(C)** 90% folded, **(D)** 85% folded, **(E)** 80% folded, and **(F)** 75% folded ubiquitin ensembles are represented using ENCORE RMSD matrix. Dimensionality reduction to 2D was performed using principal component analysis (PCA).

**Fig.S6:**
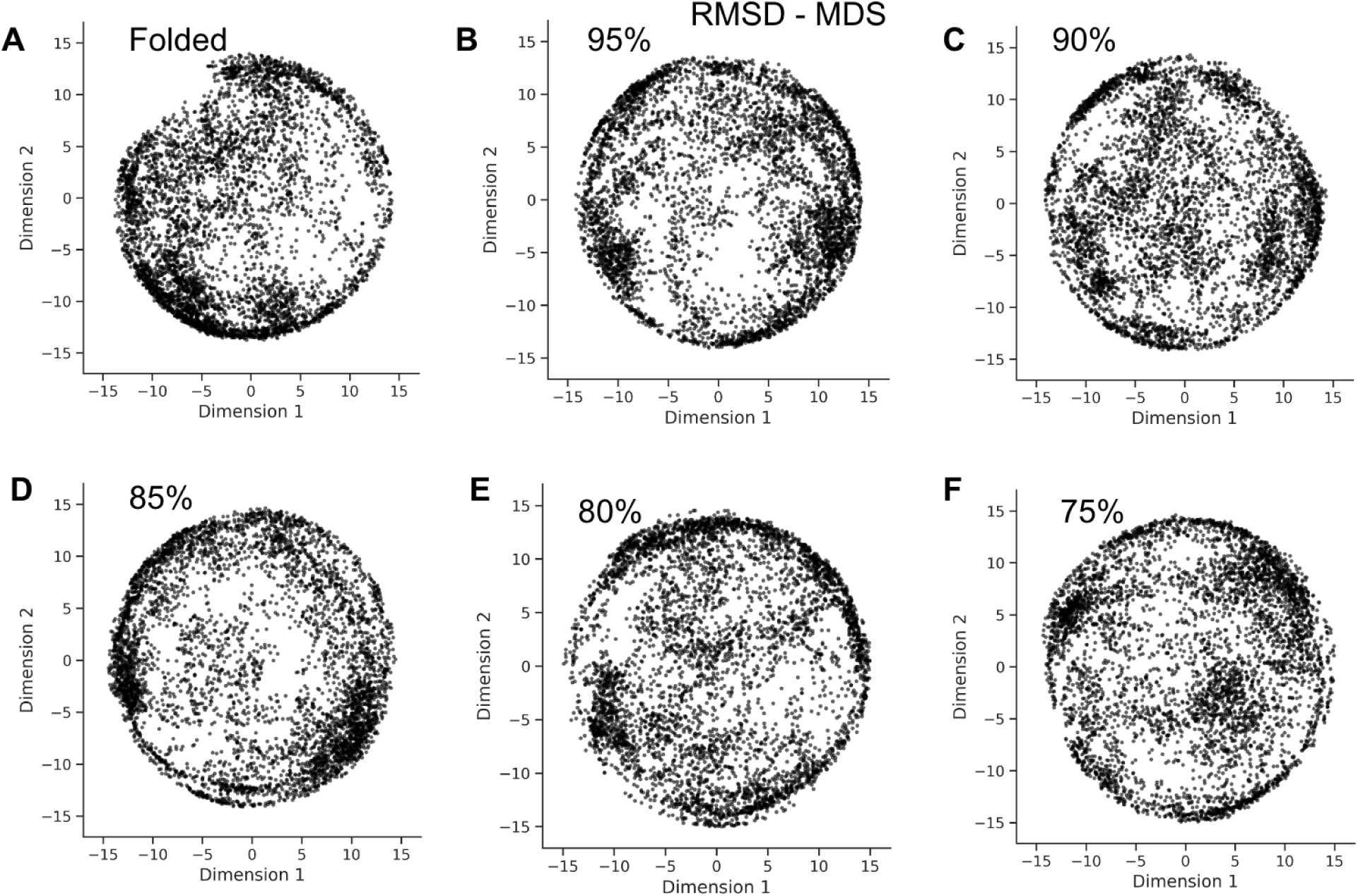
Representation and dimensionality reduction of ubiquitin ensembles. **(A)** The folded and partially folded **(B)** 95% folded, **(C)** 90% folded, **(D)** 85% folded, **(E)** 80% folded, and **(F)** 75% folded ubiquitin ensembles are represented using ENCORE RMSD matrix. Dimensionality reduction to 2D was performed using multi-dimensional scaling (MDS).

**Fig.S7:**
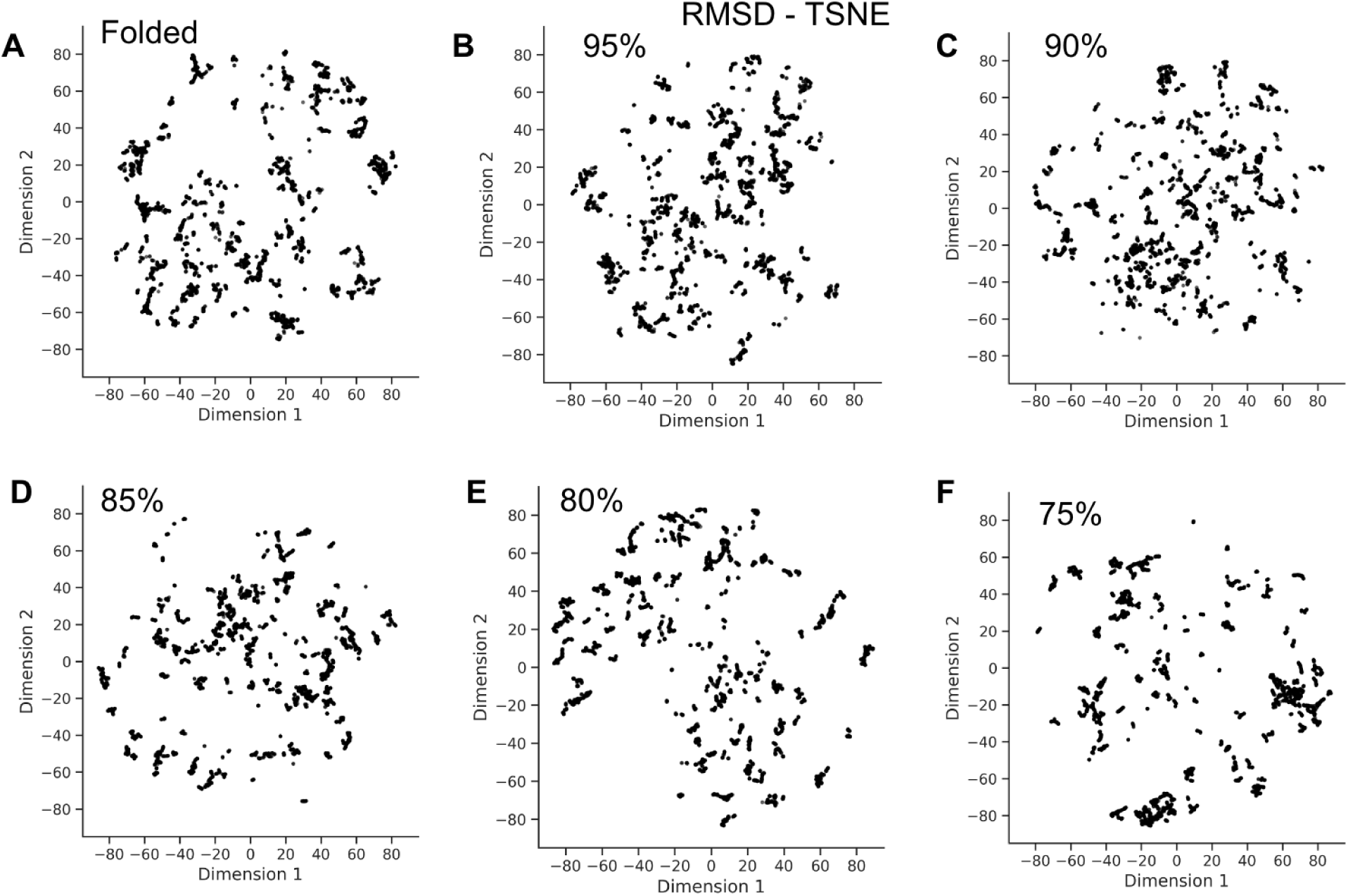
Representation and dimensionality reduction of ubiquitin ensembles. **(A)** The folded and partially folded **(B)** 95% folded, **(C)** 90% folded, **(D)** 85% folded, **(E)** 80% folded, and **(F)** 75% folded ubiquitin ensembles are represented using ENCORE RMSD matrix. Dimensionality reduction to 2D was performed using t-distributed stochastic neighbor embedding (t-SNE).

**Fig.S8:**
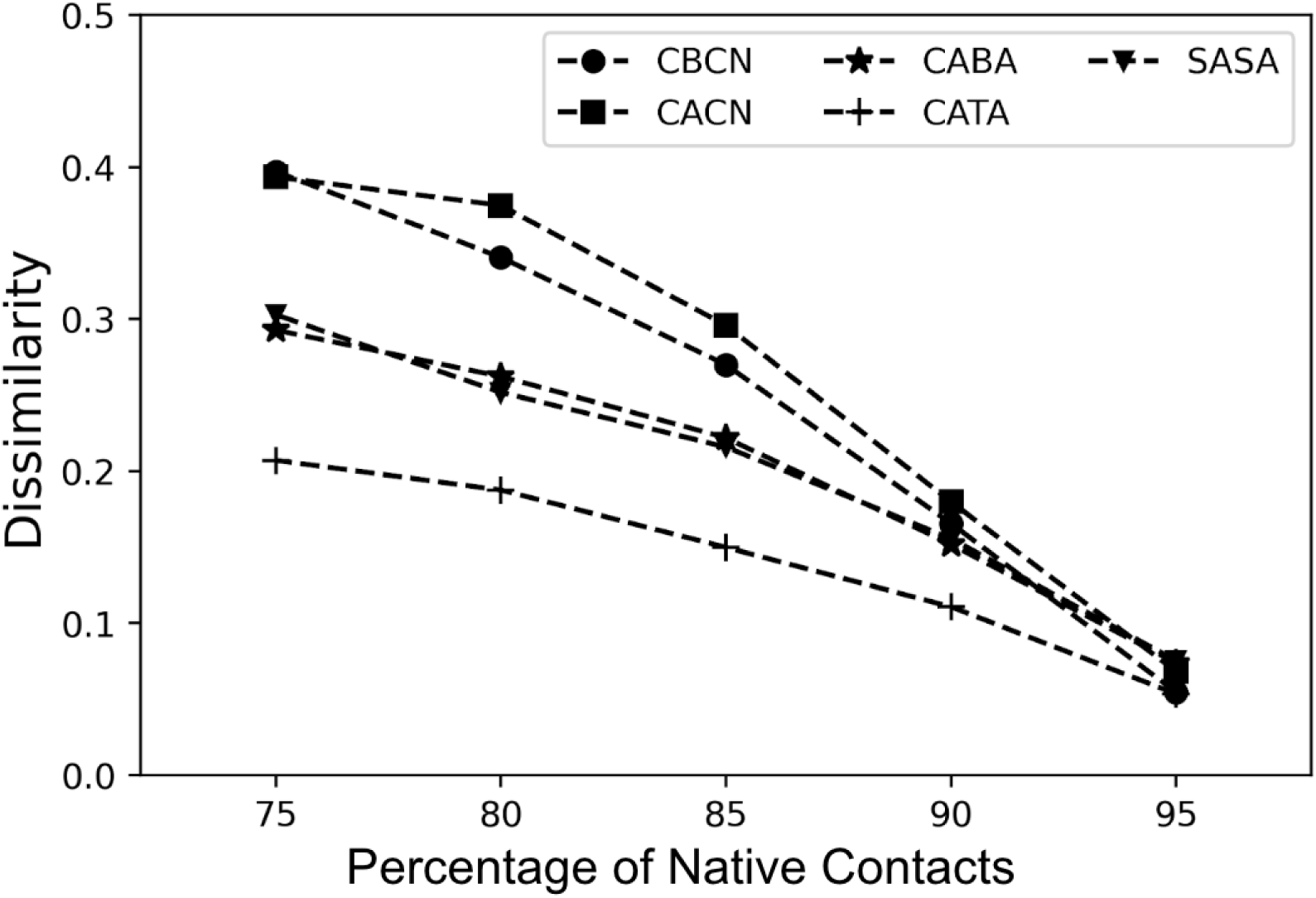
Comparing partially folded ensembles of ubiquitin. Dissimilarity in the PROTHON method (Eq. (9), main text) between partially folded ensembles (75% *< Q <* 95%) and the folded equilibrium ensemble of ubiquitin. Dissimilarity is quantified using several local structural properties, including C*_β_* contact number (circles), C*_α_* contact number (squares), virtual C*_α_*-C*_α_* bond angle, CABA (stars), C*_α_* torsion angle CATA (crosses); and per residue solvent accessible surface area, SASA (triangles).

## TOC Graphic

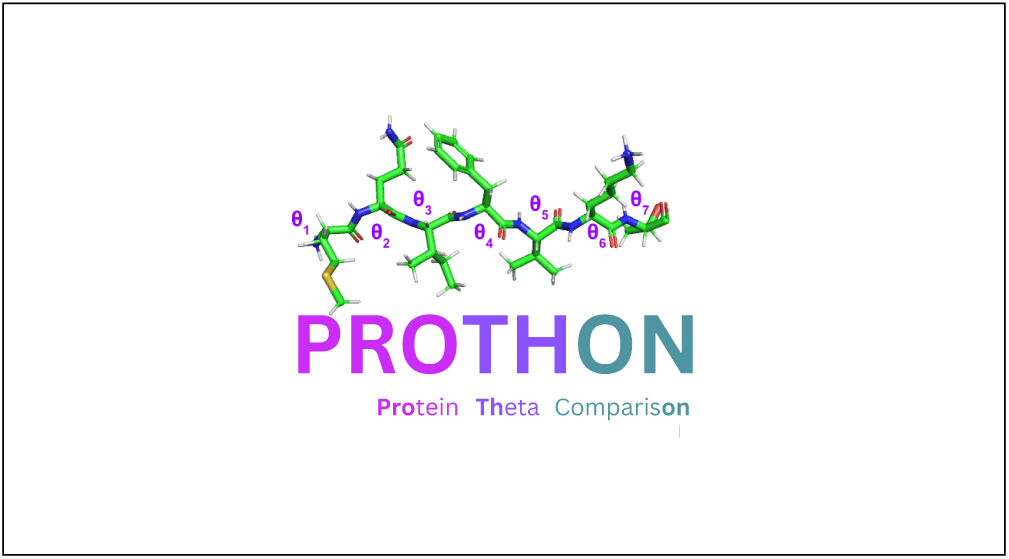

## References

(1) Goldsmith-Fischman, S.; Honig, B. Structural genomics: Computational methods for structure analysis. Protein Sci. 2003, 12, 1813–1821.

(2) Redfern, O. C.; Dessailly, B.; Orengo, C. A. Exploring the structure and function paradigm. Curr. Opin. Struct. Biol. 2008, 18, 394–402.

(3) Worth, C. L.; Gong, S.; Blundell, T. L. Structural and functional constraints in the evolution of protein families. Nat. Rev. Mol. Cell Biol. 2009, 10, 709–720.

(4) Aloy, P.; Russell, R. B. Structural systems biology: modelling protein interactions. Nat. Rev. Mol. Cell Biol. 2006, 7, 188–197.

(5) Hillisch, A. Protein structure-based drug design: applications, limitations and future developments. Chem. Cent. J. 2008, 2, S15.

(6) Zhou, W.; S^̌^midlehner, T.; Jerala, R. Synthetic biology principles for the design of protein with novel structures and functions. FEBS Lett. 2020, 594, 2199–2212.

(7) Murray, D.; Petrey, D.; Honig, B. Integrating 3D structural information into systems biology. J. Biol. Chem. 2021, 296, 100562.

(8) Staker, B. L.; Buchko, G. W.; Myler, P. J. Recent contributions of structure-based drug design to the development of antibacterial compounds. Curr. Opin. Microbiol. 2015, 27, 133–138.

(9) Śledź, P.; Caflisch, A. Protein structure-based drug design: from docking to molecular dynamics. Curr. Opin. Struct. Biol. 2018, 48, 93–102.

(10) Mohazab, A. R.; Plotkin, S. S. Structural alignment using the generalized Euclidean distance between conformations. Int. J. Quantum Chem. 2009, 109, 3217–3228.

(11) Faŕıas-Rico, J. A.; Schmidt, S.; Höcker, B. Evolutionary relationship of two ancient protein superfolds. Nat. Chem. Biol. 2014, 10, 710–715.

(12) Andersen, J. N.; Mortensen, O. H.; Peters, G. H.; Drake, P. G.; Iversen, L. F.; Olsen, O. H.; Jansen, P. G.; Andersen, H. S.; Tonks, N. K.; Møller, N. P. H. Structural and Evolutionary Relationships among Protein Tyrosine Phosphatase Domains. Mol. Cell. Biol. 2001, 21, 7117–7136.

(13) Kufareva, I.; Abagyan, R. Methods of Protein Structure Comparison; 2011; pp 231–257.

(14) Zhang, Y.; Skolnick, J. Scoring function for automated assessment of protein structure template quality. Proteins: Struct., Funct., Bioinf. 2004, 57, 702–710.

(15) Kryshtafovych, A.; Schwede, T.; Topf, M.; Fidelis, K.; Moult, J. Critical assessment of methods of protein structure prediction (CASP)—Round XIII. Proteins: Struct., Funct., Bioinf. 2019, 87, 1011–1020.

(16) Pereira, J.; Simpkin, A. J.; Hartmann, M. D.; Rigden, D. J.; Keegan, R. M.; Lupas, A. N. High-accuracy protein structure prediction in CASP14. Proteins: Struct., Funct., Bioinf. 2021, 89, 1687–1699.

(17) Plotkin, S. S. Generalization of distance to higher dimensional objects. Proc. Natl. Acad. Sci. U. S. A. 2007, 104, 14899–14904.

(18) Das, A.; Sin, B. K.; Mohazab, A. R.; Plotkin, S. S. Unfolded protein ensembles, folding trajectories, and refolding rate prediction. J. Chem. Phys. 2013, 139, 1–15.

(19) Uversky, V. N.; Gillespie, J. R.; Fink, A. L. Why are “natively unfolded” proteins unstructured under physiologic conditions? Proteins: Struct., Funct., Bioinf. 2000, 41, 415–427.

(20) Uversky, V. N. Natively unfolded proteins: A point where biology waits for physics. Protein Sci. 2002, 11, 739–756.

(21) Dunker, A. K.; Oldfield, C. J.; Meng, J.; Romero, P.; Yang, J. Y.; Chen, J.; Vacic, V.; Obradovic, Z.; Uversky, V. N. The unfoldomics decade: an update on intrinsically disordered proteins. BMC Genomics 2008, 9, S1.

(22) Ozenne, V.; Bauer, F.; Salmon, L.; Huang, J.-r.; Jensen, M. R.; Segard, S.; Bernado, P.; Charavay, C.; Blackledge, M. Flexible-meccano: a tool for the generation of explicit ensemble descriptions of intrinsically disordered proteins and their associated experimental observables. Bioinformatics 2012, 28, 1463–1470.

(23) Hsueh, S. C. C.; Aina, A.; Roman, A. Y.; Cashman, N. R.; Peng, X.; Plotkin, S. S. Optimizing Epitope Conformational Ensembles Using *α*-Synuclein Cyclic Peptide “Glycindel” Scaffolds: A Customized Immunogen Method for Generating Oligomer-Selective Antibodies for Parkinson’s Disease. ACS Chemical Neuroscience 2022, https://doi.org/10.1021/acschemneuro.1c00567.

(24) Hsueh, S. C.; Aina, A.; Plotkin, S. S. Ensemble Generation for Linear and Cyclic Peptides Using a Reservoir Replica Exchange Molecular Dynamics Implementation in GROMACS. The Journal of Physical Chemistry B 2022, 126, 10384–10399, PMID: 36410027.

(25) Brüschweiler, R. Efficient RMSD measures for the comparison of two molecular ensembles. Proteins: Struct., Funct., Bioinf. 2002, 50, 26–34.

(26) Lindorff-Larsen, K.; Ferkinghoff-Borg, J. Similarity Measures for Protein Ensembles. PLoS ONE 2009, 4, 1–13.

(27) Kullback, S.; Leibler, R. A. On Information and Sufficiency. Ann. Math. Stat. 1951, 22, 79–86.

(28) Tiberti, M.; Papaleo, E.; Bengtsen, T.; Boomsma, W.; Lindorff-Larsen, K. ENCORE: Software for Quantitative Ensemble Comparison. PLoS Comput. Biol. 2015, 11, 1–16.

(29) Koehl, P. Protein structure similarities. Curr. Opin. Struct. Biol. 2001, 11, 348–353.

(30) Lazar, T.; Guharoy, M.; Vranken, W.; Rauscher, S.; Wodak, S. J.; Tompa, P. Distance-Based Metrics for Comparing Conformational Ensembles of Intrinsically Disordered Proteins. Biophys. J. 2020, 118, 2952–2965.

(31) McClendon, C. L.; Hua, L.; Barreiro, G.; Jacobson, M. P. Comparing Conformational Ensembles Using the Kullback–Leibler Divergence Expansion. J. Chem. Theory Comput. 2012, 8, 2115–2126.

(32) Shrake, A.; Rupley, J. Environment and exposure to solvent of protein atoms. Lysozyme and insulin. J. Mol. Biol. 1973, 79, 351–371.

(33) McGibbon, R.; Beauchamp, K.; Harrigan, M.; Klein, C.; Swails, J.; Herńandez, C.; Schwantes, C.; Wang, L.-P.; Lane, T.; Pande, V. MDTraj: A Modern Open Library for the Analysis of Molecular Dynamics Trajectories. Biophys. J. 2015, 109, 1528–1532.

(34) Hinsen, K.; Hu, S.; Kneller, G. R.; Niemi, A. J. A comparison of reduced coordinate sets for describing protein structure. J. Chem. Phys. 2013, 139, 1–9.

(35) Kinjo, A. R.; Horimoto, K.; Nishikawa, K. Predicting absolute contact numbers of native protein structure from amino acid sequence. Proteins: Struct., Funct., Bioinf. 2004, 58, 158–165.

(36) Yuan, Z. Better prediction of protein contact number using a support vector regression analysis of amino acid sequence. BMC Bioinf. 2005, 6, 248.

(37) Silverman, B. W. Density Estimation for Statistics and Data Analysis. Monographs on Statistics and Applied Probability, Chapman and Hall, London 1986, 26.

(38) Virtanen, P. et al. SciPy 1.0: fundamental algorithms for scientific computing in Python. Nat. Methods 2020, 17, 261–272.

(39) Mann, H. B.; Whitney, D. R. On a Test of Whether one of Two Random Variables is Stochastically Larger than the Other. Ann. Math. Stat. 1947, 18, 50–60.

(40) Fay, M. P.; Proschan, M. A. Wilcoxon-Mann-Whitney or t-test? On assumptions for hypothesis tests and multiple interpretations of decision rules. Statistics Surveys 2010, 4.

(41) Goldstein, G.; Scheid, M.; Hammerling, U.; Schlesinger, D. H.; Niall, H. D.; Boyse, E. A. Isolation of a polypeptide that has lymphocyte-differentiating properties and is probably represented universally in living cells. Proc. Natl. Acad. Sci. U. S. A. 1975, 72, 11–15.

(42) Vijay-kumar, S.; Bugg, C. E.; Cook, W. J. Structure of ubiquitin refined at 1.8 Å resolution. J. Mol. Biol. 1987, 194, 531–544.

(43) Huang, J.; Rauscher, S.; Nawrocki, G.; Ran, T.; Feig, M.; de Groot, B. L.; Grubmüller, H.; MacKerell, A. D. CHARMM36m: an improved force field for folded and intrinsically disordered proteins. Nat. Methods 2017, 14, 71–73.

(44) Van Der Spoel, D.; Lindahl, E.; Hess, B.; Groenhof, G.; Mark, A. E.; Berendsen, H. J. C. GROMACS: Fast, flexible, and free. J. Comput. Chem. 2005, 26, 1701–1718.

(45) The PLUMED consortium, Promoting transparency and reproducibility in enhanced molecular simulations. Nat. Methods 2019, 16, 670–673.

(46) Best, R. B.; Hummer, G.; Eaton, W. A. Native contacts determine protein folding mechanisms in atomistic simulations. Proc. Natl. Acad. Sci. U. S. A. 2013, 110, 17874– 17879.

(47) Habibi, M.; Rottler, J.; Plotkin, S. S. As Simple As Possible, but Not Simpler: Exploring the Fidelity of Coarse-Grained Protein Models for Simulated Force Spectroscopy. PLoS Comput. Biol. 2016, 12, 1–31.

(48) Lever, J.; Krzywinski, M.; Altman, N. Principal component analysis. Nat. Methods 2017, 14, 641–642.

(49) Cox, M. A. A.; Cox, T. F. Handbook of Data Visualization; Springer Berlin Heidelberg: Berlin, Heidelberg, 2008; pp 315–347.

(50) van der Maaten, L. J. P.; Hinton, G. E. Visualizing High-Dimensional Data Using t-SNE. Journal of Machine Learning Research 2008, 9, 2579–2605.

(51) Pedregosa, F. et al. Scikit-learn: Machine Learning in Python. Journal of Machine Learning Research 2011, 12, 2825–2830.

(52) Arendt, T.; Stieler, J. T.; Holzer, M. Tau and tauopathies. Brain Res. Bull. 2016, 126, 238–292.

(53) Pedersen, J. T.; Sigurdsson, E. M. Tau immunotherapy for Alzheimer’s disease. Trends Mol. Med. 2015, 21, 394–402.

(54) Plotkin, S. S.; Cashman, N. R. Passive immunotherapies targeting A*β* and tau in Alzheimer’s disease. Neurobiol. Dis. 2020, 144, 1–26.

(55) Lazar, T., et al. PED in 2021: a major update of the protein ensemble database for intrinsically disordered proteins. Nucleic Acids Res. 2021, 49, D404–D411.

(56) Ozenne, V.; Schneider, R.; Yao, M.; Huang, J.-r.; Salmon, L.; Zweckstetter, M.; Jensen, M. R.; Blackledge, M. Mapping the Potential Energy Landscape of Intrinsically Disordered Proteins at Amino Acid Resolution. J. Am. Chem. Soc. 2012, 134, 15138–15148.

(57) Torda, A. E.; van Gunsteren, W. F. Algorithms for clustering molecular dynamics configurations. Journal of Computational Chemistry 1994, 15, 1331–1340.

(58) Zhou, T.; Caflisch, A. Distribution of Reciprocal of Interatomic Distances: A Fast Structural Metric. Journal of Chemical Theory and Computation 2012, 8, 2930–2937, PMID: 26592131.

(59) Wallin, S.; Farwer, J.; Bastolla, U. Testing similarity measures with continuous and discrete protein models. Proteins: Structure, Function, and Bioinformatics 2003, 50, 144–157.

(60) Hardin, C.; Eastwood, M. P.; Prentiss, M. C.; Luthey-Schulten, Z.; Wolynes, P. G. Associative memory Hamiltonians for structure prediction without homology: / proteins. Proceedings of the National Academy of Sciences 2003, 100, 1679–1684.

(61) Mariani, V.; Biasini, M.; Barbato, A.; Schwede, T. lDDT: a local superposition-free score for comparing protein structures and models using distance difference tests. Bioinformatics 2013, 29, 2722–2728.

(62) Zemla, A.; Venclovas,; Moult, J.; Fidelis, K. Processing and analysis of CASP3 protein structure predictions. Proteins: Structure, Function, and Bioinformatics 1999, 37, 22–29.

